# PSEA: A phenotypic similarity ensemble approach for prioritizes candidate genes to aid mendelian disease diagnosis

**DOI:** 10.1101/2021.10.13.464308

**Authors:** Zhonghua Wang, Lipei Liu, Chao Chen, Xi Liu, Fei Tang, Yixiang Zhang, Yunmei Chen, Yaoshen Wang, Jun Sun, Zhiyu Peng

**Author notes:** These authors contributed equally to this work (Zhonghua Wang, Lipei Liu, Chao Chen).

## Abstract

**Motivation:** Next-generation sequencing (NGS) is increasingly applied to the molecular diagnosis of genetic disorders. However, challenges for the interpretation of NGS data remain given the massive number of variants produced by NGS. Careful assessment is required to identify the most likely disease-causing variants that best match the patients’ clinical phenotypes, which is highly experience-dependent and of low cost-effectiveness.

**Results:** The human phenotype ontology (HPO) together with the information content (IC) are widely used for phenotypic similarity evaluation. Here, we introduce PSEA, a new phenotypic similarity evaluation tool capable of quantifying groups of HPO terms unbiasedly. By comparing with other methods, PSEA show optimal performance and show a higher tolerance to phenotypic noise or incompleteness. We also developed a web server for disease-causing gene prioritization and HPO-gene weighted linkage visualization.

**Availability:** Source code and Web service are free available at https://github.com/zhonghua-wang/psea and https://phoenix.bgi.com/psea, respectively.

**Supplementary information:** Supplementary data are available at *Bioinformatics* online.

## 1 Introduction

Phenotypic similarity evaluation plays an essential role in Mendelian disorder diagnosis (Chen *et al*., 2019). Many computational approaches were developed to prioritize candidate genes and aid the interpretation of next-generation sequencing data. Generally, these methods encompass two lines of strategies, i.e., genotype-driven, and phenotype-driven. Genotype-driven methods use sequence and genomic attributes such as variation type, allele frequency, interspecies conservation, and other in silico prediction data to rank candidate genes (Kircher *et al*., 2014). Unlike genotypic data, phenotypic data is less standardized. Some clinical phenotypic information is still presented as a mixture of text, thus difficult to be automatically processed (Wei and Denny, 2015). There are several phenotype-driven approaches that prioritize and predict candidate genes. Xrare (Li *et al*., 2019) is a comprehensive method which utilizes a phenotype-driven similarity scoring system called Emission-Reception Information Content (ERIC) and a genotype system to build a random forest model for pathogenicity prediction. Exomiser utilizes genotype-phenotype associations in the model organisms (Robinson *et al*., 2014). Phenotype risk score (PheRS) (Clark *et al*., 2019) translates HPO terms to phecodes and assigns risk scores to clinical presentations, thus estimate pathogenicity of rare variants, and predict causal genes.

In the present report, we describe PSEA (Phenotypic Similarity Ensemble Approach), a new bioinformatic method for phenotype-driven gene prioritization. PSEA utilizes phenotype-gene correlations in the Human Phenotype Ontology (HPO)(Köhler *et al*., 2021) database and assigns similarity scores between clinical symptoms and presentations of a candidate gene based on the information content (IC) of corresponding HPO terms. We conducted an intense data simulation to balance the impact of variable HPO numbers associated with a gene and normalized similarity scores to Z-scores. Comparing with other popular prioritizers, PSEA showed superior accuracy and a better ability to counteract phenotypic noise.

## 2 Implementation

HPO, which is a standardized vocabulary of human phenotypic abnormalities, is widely used in rare disease diagnosis. The information content (IC) or IC-derived value of HPO term was calculated for phenotypic similarity evaluation. ERIC (Emission-Reception Information Content) score, an IC-derived method introduced by Li et al (Li *et al*., 2019). which is highly tolerant of incomplete and imprecision in clinical phenotypes, were proved to be a promising measurement in gene prioritization. This method was employed to calculate the raw similarity between two HPO terms.

**Figure 1.**
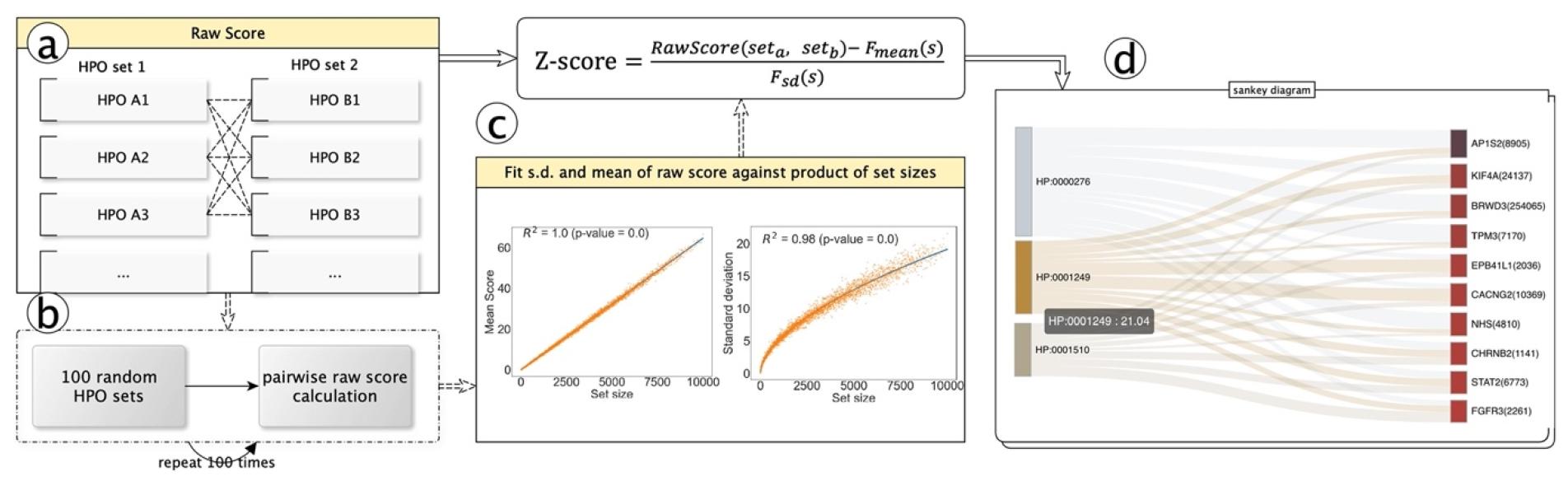
Schematic overview of PSEA. (a) Raw score of two HPO sets is calculated by the sum of pairwise similarity of HPO terms. (b) Random HPO sets are created. (c) Statistic model is built, and the raw score is normalized to z-score. (d) Connections between prioritized genes and HPO terms are illustrated in a sankey diagram.

As the annotated HPO number of a gene vary widely, for example, global development delay (HP:0001263) is currently annotated by 1,442 genes, while other HPO terms may have little to no genes associated, the likelihood of two HPO sets will grow with the size of the sets if using the sum of raw score as the measurement. To minimize the influence of HPO size, we introduce the Z-score to normalize the score. The statistical models of standard deviation and expect score of given set size were determined/fitted by 495,000 pairs of HPO sets, which were randomly populated from the HPO database with set size range from 1 to 100. The detail procedure of building PSEA model can be found in supplementary materials. The normalized Z-score endow the standardization characteristics to PSEA, which make the scores of different HPO set pairs are comparable.

We developed an interactive web server and intergraded doc2hpo (Liu *et al*., 2019) which can extract HPO terms from clinical description automatically, for user to select disease-causing genes online easily. Moreover, to better illustrate how each HPO term contributed to final score, a Sankey diagram was provided by the PSEA web server, which can help visualizing the weightiness and specificity of given HPO terms.

## 3 Results

To demonstrate the performance of PSEA, we retrospectively analyzed 420 in-house genetic cases. These patients were tested by Exome sequencing with causal variants identified and clinical phenotypes were recorded as HPO terms. Sequencing variants were interpreted according to the joint guideline of the American College of Medical Genetics and Genomics and the Association for Molecular Pathology (Richards *et al*., 2015). PSEA outperform other methods with ranked 21.31% (vs. 18.83%), 39.79% (vs. 35.52%), 61.28% (vs. 55.06%), 70.87% (vs. 68.56%), and 82.6% (vs. 80.46%) genes at top 1, top 3, top 10, top 20, and top 50, respectively. Considering the imprecise and noisy nature of clinical phenotypes, 1) original HPO terms were replaced with their parent HPO terms, sibling HPO terms or random HPO terms to mimic phenotypic imprecision; 2) half of the original HPO terms were randomly discarded to mimic phenotypic incompleteness; 3) parent or sibling HPO terms were added to original HPOs to mimic phenotypic redundancy. In all circumstances, PSEA also shown best ranking performance (see Supplementary Material).

In summary, we have developed a novel method that addresses the need for more accurate and robust phenotypic similarity measurement. This method shown better prioritizing performance and higher tolerance to phenotypic noise or incompleteness. We expect that the robust and standardization characteristics of PSEA will help researchers and clinicians quick and better identify disease-causing gene or variants.

## Acknowledgements

We thank all the users of PSEA for their valuable and constructive feedbacks.

