## Supplemental material for "PSEA: A phenotypic similarity ensemble approach for prioritizes candidate genes to aid mendelian disease diagnosis"

### Method

HPO, which is a standardized vocabulary of human phenotypic abnormalities, is widely used in rare disease diagnosis. The information content (IC) or IC-derived value of HPO term was calculated for phenotypic similarity evaluation. ERIC (Emission-Reception Information Content) score, an IC-derived method introduced by Li et al (Li *et al.*, 2019). which is highly tolerant of incomplete and imprecision in clinical phenotypes, were proved to be a promising measurement in gene prioritization. We used this method to calculate the similarity between two HPO terms. The procedures for building the PSEA model are described as follows.

1. Similarity measurement between two HPO terms

Firstly, the IC of an HPO term was computed based on the HPO database (HPO database of accessed date 2020-03-13 was used in this study), and then it was converted to the ERIC score (eq 1, eq 2). It's worth to mention that there are two different gene-phenotype mapping file on HPO website; 1) genes to phenotype mapping file, it provides a link between genes and HPO terms that has most specificity, while 2) in phenotype to genes mapping file, it also mapped genes to the ancestor classes of each associated phenotype (please refer to https://hpo.jax.org/app/faq for detail description). We used the later one ,which is more informative, for PSEA modeling.

$IC(hpo)=-log(n/N)$ … (eq 1)

where n stands for the number of genes associated with the current HPO and N is the number of total genes under consideration.

$SIM_{H}\left( hpo_{a},hpo_{b} \right)=Max\left( 0,2*IC\left( t_{M} \right)-Min\left( IC\left( hpo_{a} \right),IC\left( hpo_{b} \right) \right) \right)$	… (eq 2)

in which t_M_ represents the most informative common ancestor, the term with maximum IC value, of hpo_a_ and hpo_b._

2. Similarity measurement between two HPO sets

As the annotated HPO number of a gene vary widely, for example, global development delay (HP:0001263) is currently annotated by 1,442 genes, while other HPO terms may have little to no genes associated, the likelihood of two HPO sets will grow with the size of the sets if using the raw score as the measurement. To minimize the influence of HPO size, we introduce the Z-score to normalize the raw score. The background data set were randomly drawn from the HPO database with set sizes ranging from 1 to 100 and an interval step of 1, where results in 4,950 pairs of HPO sets. Then the pairwise raw score of HPO sets was calculated. This procedure was repeated 100 times.

The mean and the SD (standard deviation) of the raw score were fitted linearity and nonlinearity against the product of the set sizes, respectively. Both fits were determined with the SciPy linear least-squares optimizer. The final set comparison Z-score was calculated as a function of the set rawscore, expected rawscore and SD (see eq 4).

$RawScore(gene_{a},gene_{b})=\sum_{i} \sum_{j} (SIM_{H}(i,j))$	… (eq3)

where i and j are the number of HPO terms within gene_a_ and gene_b_ respectively.

$\text{Z-score}=\frac{{RawScore\left( {set}_{a}{, set}_{b} \right)- F}_{mean}(s)}{F_{sd}(s)}$ … (eq4)

Where s is the product of HPO set a and b, F_mean_ and F_sd_ are:

$F_{mean}(x)= \mu x$	…. (eq5)

$F_{sd}\left( x \right) = \varphi x^{\varepsilon}$	… (eq6)

Where the parameters μ, ϕ and ε were determined by fitting the random background statistical model.

### Performance evaluation on WES data sets

To evaluate the performance of PSEA, we retrospectively analyzed 420 genetic cases, including 563 variants from 358 genes, that went through exon sequencing (ES) in BGI from 2019 to 2020. These patients were tested by ES with causal variants identified and clinical phenotypes recorded as HPO terms.

Sequencing variants were interpreted according to the joint guideline of the American College of Medical Genetics and Genomics and the Association for Molecular Pathology (Richards *et al.*, 2015). Briefly, we assigned pathogenicity to the variants based on: 1) the minor allele frequency ≤1% in the databases of ExAC (http://exac.broadinstitute.org) and gnomAD (https://gnomad. broadinstitute.org); 2) the inheritance pattern was consistent with family segregation information; 3) supportive evidence from literature and in-house database; 4) null variants in a gene when LOF is a known mechanism of disease; 5) conservation and predicted impact on coding and noncoding sequence; 6) relevance to the current phenotype.

Free-text clinical presentations were manually extracted as HPO terms by clinical experts. To study the phenotype-driven gene prioritization performance of PSEA, we did not incorporate genotypic information of the variants. Only genes deemed relevant to the clinical presentations went through further analysis.

For each case, HPO-coded phenotypic information was inputted and a prioritized list was then obtained. For each method, we recorded the ranks of causal genes presented in corresponding lists and collectively calculated the cumulative frequency (CF) of consecutive ranks. To independently test the phenotypic prioritization ability, genotype-derived results were discarded. Therefore, a higher CF value stands for better accuracy. In the present study, three public tools were tested in parallel on the same ES dataset. There tools were Xrare (docker-image xrare-pub:2015), Exomiser (version v11), and PheRS (phecode version v1.2).

All patients of these data sets signed informed consent and allowed their data to be reanalyzed and published for a research purpose without personally identifiable information. The Institutional Review Board (IRB) of BGI approved this study.

### Construction of phenotypic noise test set

In this study, clinical information was standardized as a set of HPO terms. To evaluate the tolerant ability of PSEA to phenotypic noise, we simulated three groups of phenotypic data based on original manually-extracted HPO terms, including imprecision, incompleteness, and redundancy groups. Briefly:

1) imprecision : we replaced original HPO terms with their parent/sibling HPO terms or added noise HPO terms to emulate imprecise phenotypic information. For original HPO terms associated with more than one sibling HPO terms, we randomly selected half of their siblings. When adding noise HPO terms to the original HPO set, 1 to 5 HPO terms unrelated to the current presentations were randomly inputted as noise.

2) incompleteness: to simulate incomplete phenotypic information, half of the original HPOs were discarded.

3) redundancy: to simulate redundant phenotypic information, we added parent or sibling HPO terms to the original HPO sets. For original HPO terms associated with more than one sibling, half of their sibling were randomly selected.

### PSEA demonstrated optimal ability in gene prioritization

We compared PSEA with other methods including ERIC of Xrare, EXO, and PheRS. Among all methods tested, PSEA demonstrated optimal ability in gene prioritization (**Figure S1**). PSEA placed 21.31% (120/563) genes at the top of the list and demonstrated superior performance over ERIC (18.83% at top 1, 106/563), EXO (15.99% at top 1, 90/563), and PheRS (9.24% at top 1, 52/563). Additionally, PSEA ranked 61.28% genes in Top 10 ranks compared to 55.06% for ERIC, 50.80% for EXO, and 30.73% for PheRS. The median rank of causal genes predicted by PSEA was 6, which was significantly better than that of ERIC (8, *p*<0.001), EXO (10, *p*<0.001), and PheRS (33, *p*<0.001).

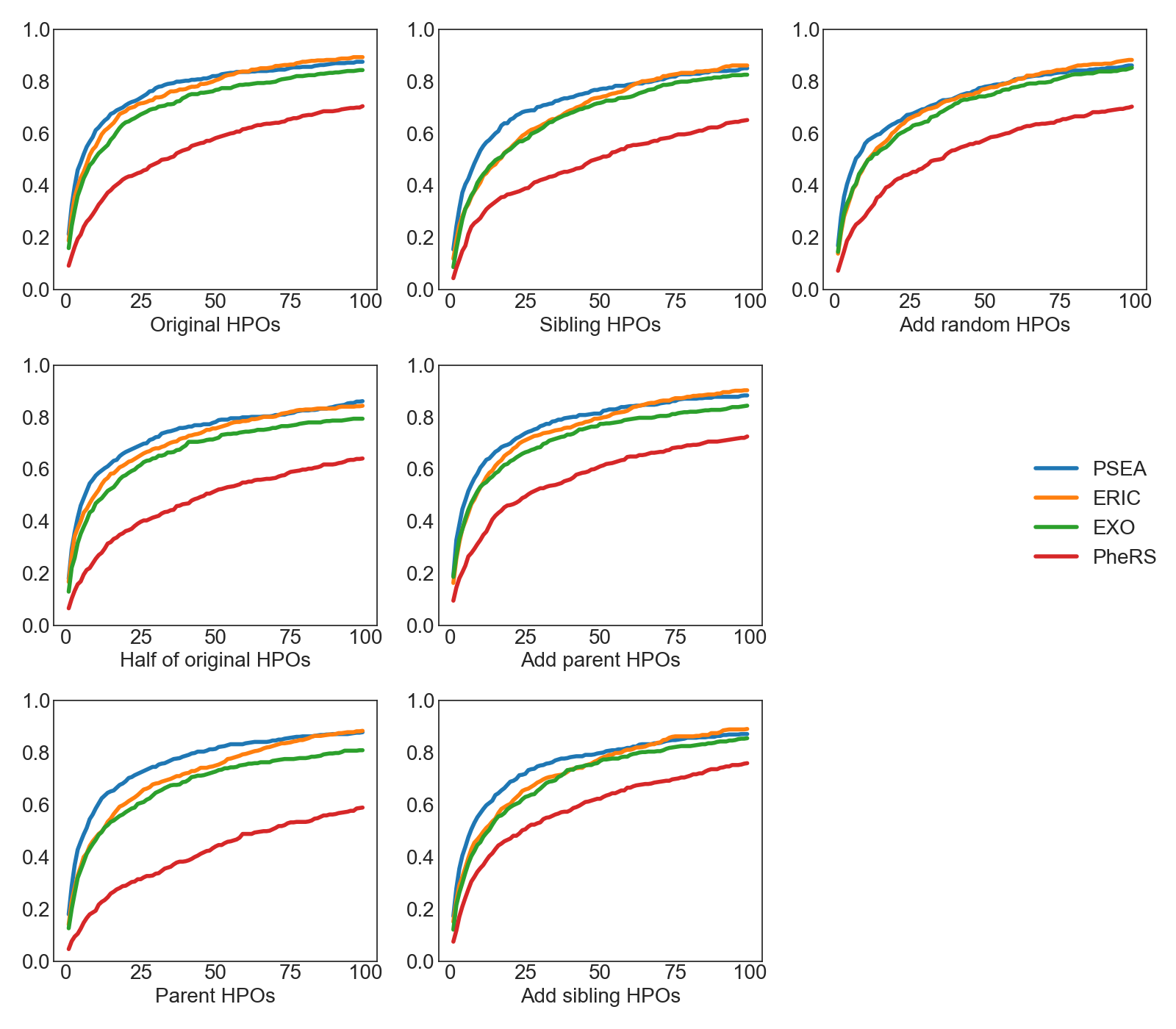

Figure S1. Performance comparison of different methods with original HPOs and 6 simulated HPOs. The x-axis is the rank percentile of target genes. The y-axis is the cumulative distribution function of the rank percentile.

We then retrieved causal genes successfully predicted by each method (defined as ranking in Top 10 by each method), thus four groups of genes were obtained (**Figure S2**). In total, 400 genes were successfully predicted, among which 120 genes were simultaneously predicted by all four methods. PSEA outperformed other methods and contributed 345 (86.25%) confirmed causal genes in Top 10 [compared to 310 genes (77.5%) by ERIC, 286 genes (70.5%) by Exomiser, and 173 genes (43.25%) by PheRS]. Meanwhile, PSEA achieved 37 unique predictions (9.25% of its total prediction), whereas ERIC was 15 (3.75%), Exomiser was 18 (4.5%), and PheRS was 8 (2%).

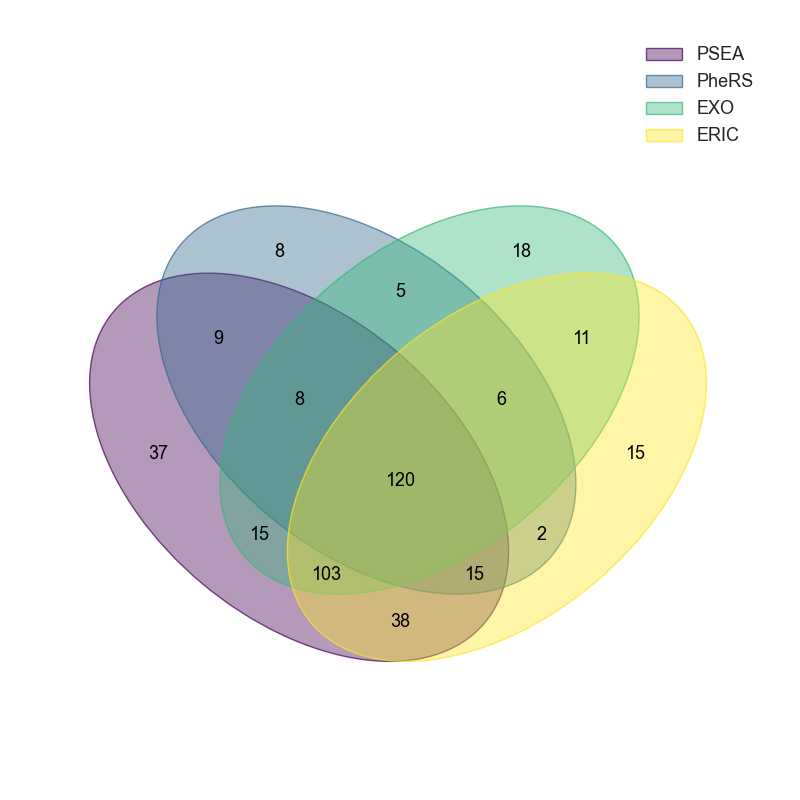

Figure S2. The shared prediction of the four different methods at top 10.

### Performance evaluation of PSEA with noise phenotypic information

The heterogeneous nature of phenotypic information remains a substantial challenge to the diagnosis of Mendelian diseases. To unveil the detailed impact of phenotypic noise on these bioinformatic tools, we challenged PSEA and other tools with imprecise, incomplete, and redundant phenotypic information. Briefly, original HPO terms (Original-HPOs) were replaced with their parent HPO terms (Replace-parent-HPOs) or sibling HPO terms (Replace-sibling-HPOs) or supplemented with random HPO terms (Add-random-HPOs) to emulate phenotypic imprecision. Half of the original HPO terms (Half-of-original-HPOs) were randomly discarded to emulate phenotypic incompleteness. Parent or sibling HPO terms were added to Original-HPOs to emulate phenotypic redundancy.

We simultaneously treated PSEA with original clinical information (Original-HPOs), imprecise (Replace-parent-HPOs/Replace-sibling-HPOs/Add-random-HPOs), incomplete (Half-of-original-HPOs), and redundant (Add-parent-HPOs/Add-sibling-HPOs) clinical information. Our simulation indicates that providing PSEA with original HPOs demonstrated the most accurate prediction of causal genes. Whereas introducing phenotypic noise considerably compromised PSEA’s performance (**Supplementary Figure S3**). Generally, phenotypic redundancy (Add-parent-HPOs/Add-sibling-HPOs) had less impact comparing to imprecise (Replace-parent-HPOs/Replace-sibling-HPOs/Add-random-HPOs) and incompleteness (Half-of-original-HPOs), while replacing Original HPOs with their siblings (Replace-sibling-HPOs) appeared the worst impact. Add-parent-HPOs showed the least impact on the predicting performance of PSEA with a 2.13% reduction of the likelihood of causal gene ranking at Top 1 comparing to Original HPOs. Whereas Replace-sibling-HPOs (original HPO terms replaced with their siblings) undermined PSEA’s performance to the greatest extent with a 5.86% reduction at Top 1. PSEA demonstrated higher accuracy over other methods in all types of phenotypic noise simulated (Figure S1).

|  | **HPO** | **HPO-half** | **HPO-parent** | **HPO-sibling** | **HPO-add-parent** | **HPO-add-sibling** | **HPO-noise-median** |
| --- | --- | --- | --- | --- | --- | --- | --- |
| **ERIC** | 8 | 10 | 13 | 17 | 9 | 12 | 12 |
| **PheRS** | 33 | 47 | 68 | 49 | 26 | 24 | 35 |
| **EXO** | 10 | 13 | 13 | 16 | 9 | 13 | 12 |
| **PSEA** | 6 | 7 | 7 | 9 | 6 | 7 | 7 |

Table S1.

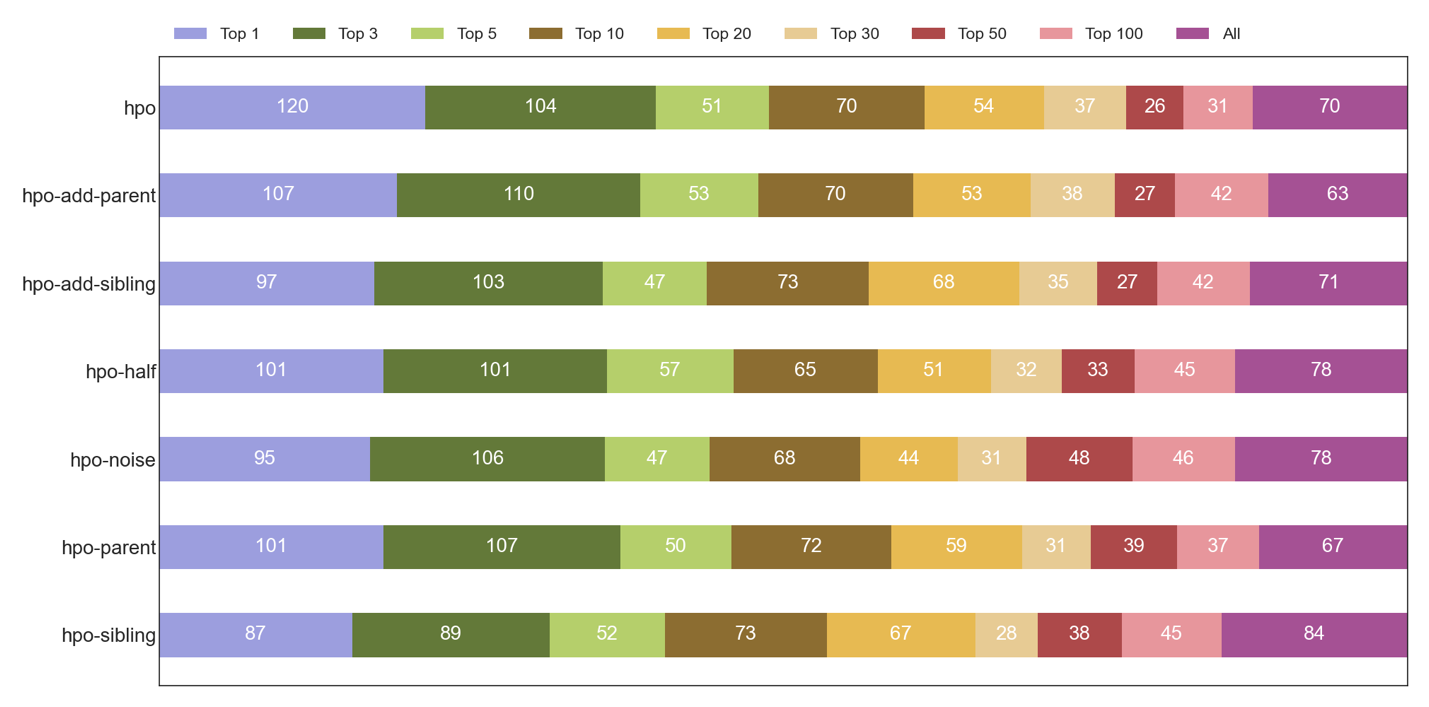
We did a statistical analysis and calculated the median rank of different methods (**Table S1**). Among all scenarios tested, PSEA demonstrated increased ability in handling phenotypic noise compared with other methods.

Figure S3. PSEA performance with different HPO noises.

### PSEA web service

Web interface of PSEA is shown in Figure S4. User can either search and select HPO terms or provide clinical description to perform gene prioritization. After providing HPOs, prediction will run automatically, and the result gene list will be shown in Sankey diagram. The width of the link between HPO and gene is directly proportional to the contribution to the similarity of selected HPO terms and the gene. User can remove or add HPO terms interactively, the Sankey diagram will be updated accordingly. The prioritized gene list as well as the Sankey diagram are both downloadable. Instructions also available online (https://phoenix.bgi.com/help#psea).

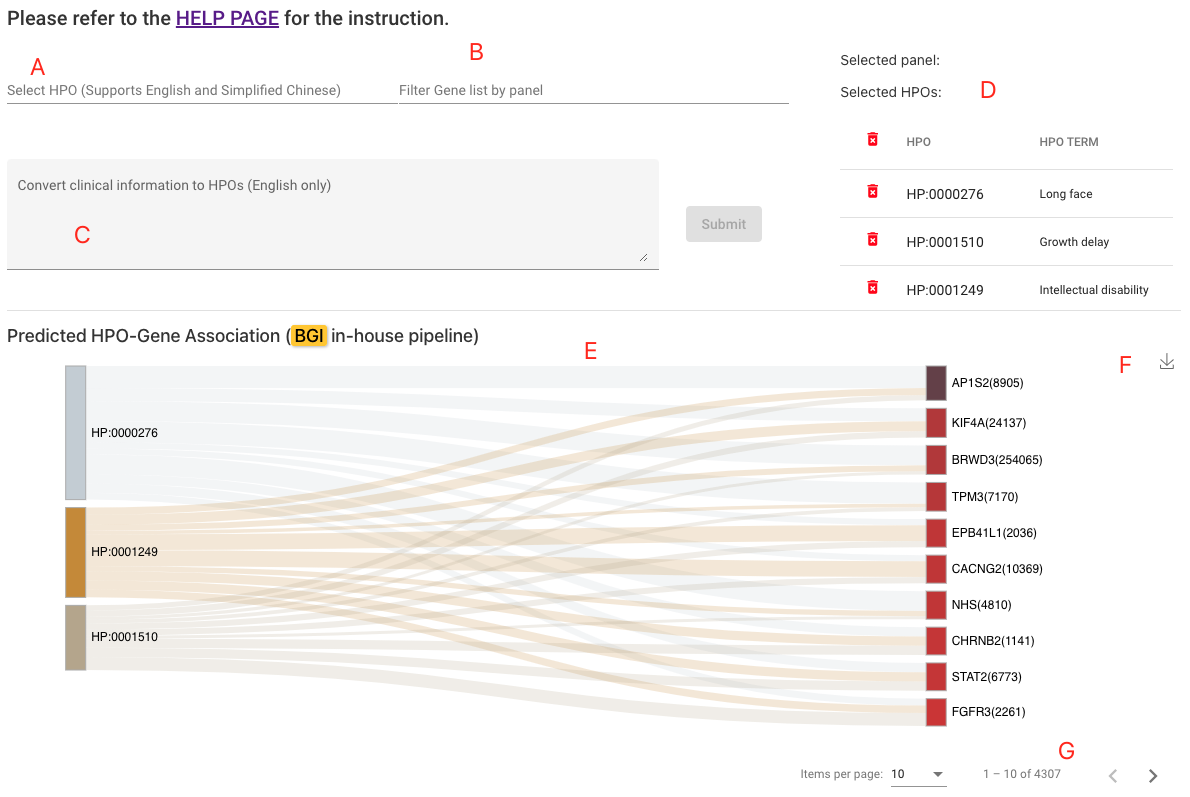

Figure S4. PSEA web page. (A) Type to search and select HPO terms, supports English and simplified Chinese. (B) Filter result with gene panels, panels were collected from Genomics England PanelApp (<https://panelapp.genomicsengland.co.uk/>). (C) Convert clinical text to HPO terms (English only), provided by doc2hpo (Liu *et al.*, 2019). (D) Manually selected HPO or automatically extracted HPO terms. (E) Sankey diagram illustrating HPO-Gene connections.
